# Prevalence and determinants of smoking status among university students: Artvin Çoruh University sample

**DOI:** 10.1101/361345

**Authors:** Dilek Karadoğan, Özgür Önal, Yalçın Kanbay

**Affiliations:** Department on Chest Diseases, Recep Tayyip Erdoğan University, School of Medicine, Rize, Turkey; Department of Public Health, Süleyman Demirel University, School of Medicine, Isparta, Turkey; Department of Psychiatric Nursing, School of Health Science, Çoruh University, Artvin, Turkey

**Keywords:** Cigarette smoking, prevalence, determinants, university students

## Abstract

**Background and aim:** While smoking prevalence has decreased in many developed countries, in others it is still an increasing risk factor for public health, especially among young adults. In this study we aimed to find the factors affecting university students’ smoking status.

**Methods:** This cross-sectional study was conducted between March and June 2017 with a simple random sampling method. A self-administered questionnaire was used to collect information on sociodemographic characteristics, cigarette smoking status, and the related risk factors. Univariate analysis and multivariate logistic regression analysis with the Backward:LR test were performed.

**Results:** Out of 2723 students, 2505 students’ data were available and suitable for analysis. Mean participant age was 20.9±2.5, with female dominance (58.9%). Students studying in two-year degree departments comprised 45.6% of the study population, and the remaining 54.4% of the students were in 4-year degree departments. In terms of parental smoking habits, 36.1% reported a smoker father, while that rate was 10.3% for mothers and 15.0% for siblings. Current smokers represented 27.9% of the group: 46% among males and 15.3% among females. Mean age for beginning smoking was 16.34±2.72, 15.65±2.67 for males, 16.34±2.72 for females (p<0.05). Mean Fagerströmtest score was 4.43±1.82, and female students had lower test scores than males (p<0.05). After controlling for potential confounders in multivariate analysis, it was seen that five factors were positively associated with current smoking (p<0.05): male gender (OR:3.43; 95%, CI:2.75-4.28), studying in two-year programs (OR: 1.74; 95% CI: 1.39-2.18), having at least one close family member who is a smoker (OR:1.63; 95% CI:1.31-2.04), having close friends who are all smokers (OR:1.81; 95% CI: 1.40-2.33), and alcohol consumption (OR:4.39; 95% CI: 3.51-5.49).

**Conclusion:** Our study showed a higher smoking rate among our study population, both compared to similar nationalstudies and the country’s overall smoking rate. Underlying factors should be evaluated via qualitative studies and preventive strategies should be implemented accordingly.

## Introduction

Cigarettes are the most commonly used form of tobacco and one of the major causes of preventable diseases globally (1). Turkey’s fight against tobacco depends on following the evidence-based tobacco control measures and policies that have been identified as effective by the World Health Organization (WHO) (2). Turkey protects its population with five tobacco control measures, known as ‘MPOWER’ policies at the highest level(3, 4). Turkey’s smoking prevalence among individuals over 15 years of age was 44.5% in 1988; after anti-tobacco policies were implemented that rate decreased to33.6% in 1993. In 2004 Turkey signed the Framework Convention on Tobacco Control (FCTC) with WHO, and by 2008 the smoking rate for that population was down slightly to 31.2% (4). A recent study explained that despite the strict anti-tobacco policies and decreased legal tobacco product sales, as of 2006 the Turkish current smoking rate had plateaued after a 20 year decline, still leaving Turkey with a higher smoking rate than other countries, such as the United States (5). According to the last Global Adult Tobacco Survey, that rate finally decreased to 27.1% in 2012; the MPOWER policies are the main factor in that result (4). Among the studies that isolated university students’ smoking rates, the situation was similar to the general population. After the implementation of MPOWER policies, researchers also found a decrease in smoking prevalence among university students from 2005-2006 to 2012-2013: 26.9% to 18.5% (6).

In general, smoking rates in Turkey differ nationwide according to socioeconomic status. A national study found no difference before and after the tobacco control policies in the poorest populations, while the smoking rate did decrease among the richest population (3). The continued levels of smoking rates in the poorest population, despite decreased legal cigarette sales, were attributed to illegal tobacco use, either as cigarettes or in other forms (3). In a recent study Turkey’s overall illicit cigarette use had been found to be around 12%, most commonly in the eastern part of the country (7). Therefore prevalence studies should take place periodically and should also evaluate regional data.

Youth are the main target for tobacco companies, and the age of starting to smoke is decreasing in developing countries. Smoking for the first time before the age of 18 has been found to be a factor contributing to life-long smoking; for those smokers quitting will be more difficult (8). Universities are an important reflection of the general data on the younger population’s smoking behaviors. Because in Turkey university students traditionally leave their homes to move to another city and start living with their peers, family pressure and control decrease and students feel much freer to make their own decisions. Therefore university life itself can became a risk factor for both initiation of and increase in smoking behavior. Accordingly, studies that compare first and last year university students’smoking levels (9, 10)— and studies that follow Turkish students prospectively—showed an increased smoking rate during university life (11). Preventions during this period are important, such as campus-wide smoking bans, educational symposiums about the harms of tobacco, and other measures that are known to negatively impact smoking rates (12, 13).

The factors associated with smoking among university students have been evaluated by previous national and international studies. The primary factors associated with student smoking status have been found to be socioeconomic status, family/friends’ smoking behavior, alcohol use, gender, faculty, year of education, and residency with friends (14, 15). However, different geographical regions have different risk factors due to their cultural-sociological differences; therefore there is a need to conduct studies to evaluate different regions’ status. Our setting for this study is a university located in the Eastern Black Sea region of Turkey. It has no smoking bans on campus and the students are mostly from the eastern and north eastern parts of the country, the children of low and middle income families. As mentioned above, those areas of the country have the highest illicit tobacco use in Turkey, both as thin paper rolled cigars and as cigarettes. Also, the city where the university is located shares a border with Georgia and is involved in that country’s illicit tobacco transport (16). This study aimed to evaluate the current smoking prevalence of these university students and the influencing factors on their current smoking status, whichwill be important to gain actual data about the eastern part of the country’s smoking trends. It may also provide data that assists in strengthening policies against smoking and ultimately aids universities to develop anti-smoking programs.

## Methods

### Settings and study design

This cross-sectional study was conducted at Çoruh University in Artvin. Artvin is a small city that is located in the Eastern Black Sea region of Turkey and has a border with the neighboring country of Georgia. This study included 9 different schools or faculties within the university—Faculty of Education, Faculty of Art and Sciences, Faculty of Economics and Administrative Sciences, Faculty of Management, Faculty of Health Sciences, Faculty of Forestry, Faculty of Engineering, Vocational School, and the vocational high school—in the academic year 2016/2017 during the period from March 1 to June 30, 2017. Çoruh University had approximately 9000 students in that education year, and at the time of data collection there were 6583 students in those departments. Based on the assumption of a margin of error of 2% and a confidence interval of 99%, the initial/minimum sample size was calculated to be 2549 students. The investigators ultimately decided to distribute the survey to 2741 students, accounting for an expected number of nonresponses and incomplete questionnaires, by using a simple random sampling method. After excluding any incomplete and unsound responses, the study sample utilized data from 2505 students.

The research was approved by an Ethical Committee and the Vice Rector of Çoruh University.

### Data collection

For the current cross-sectional study a short self-administered questionnaire was used. The questionnaire was succinct to encourage the students to respond. It comprised 26 questions that were modified from previously used surveys in Turkey (15, 17, 18) including 12 questions about the socioeconomic profile of the respondents and 8 questions about their attitude toward smoking and alcohol. The last 6 questions, the Fagerström test (19), were for the regular smokers. A smoker in this study was defined as a participant who had smoked regularly in last 30 days prior to taking the questionnaire and had smoked at least 100 cigarettes in their lifetime. A non-smoker was defined as someone who did not smoke in the previous 30 days and/or had not smoked 100 cigarettes lifetime or had smoked over 100 cigarettes lifetime but not smoking in last 30 days. A team of 10 assistants was trained to ensure a unified procedure for data collection. A preliminary pilot study was undertaken with 50 students and some questions were subsequently rephrased and modified accordingly.

### Statistical analysis

Data were entered and analyzed using Statistical Package for Social Sciences (SPSS) version 20.0. Descriptive statistics and logistic regression were performed. For categorical variables Pearson’s Chi Square test was used, for continual variables Students’ t-test was used. Multivariate logistic regression analysis with Backward LR was used to identify factors independently associated with the smoking status of university students. A p value of< 0.05 showed statistical significance.

## Results

In total 2505 students’ data were evaluated; 58.9% of them were female and 41.1% of them were male. Mean age of the students was 20.87±2.50. Overall smoking rate of the students was 27.9%: among females that rate was 15.9% and among males that was 46.0%. Detailed demographical characteristics of the study population are seen in Table 1 and Table 2.

**Table 1.** Sociodemographic characteristics of the students according to smoking status (Categorical variables).

|  |  | Smoker<br>(699, %27.9)<br>n(%row)* | Total<br>(2505, %100.0)<br>n(%column)** | p |
| --- | --- | --- | --- | --- |
| **Gender** | **Female** | 226 (15.3) | 1476 (58.9) | <0.001 |
|  | **Male** | 473 (46.0) | 1029 (41.1) |  |
| **Consumption of alcohol** | **Present** | 343 (59.7) | 575 (23.0) | <0.001 |
|  | **Absent** | 356 (18.4) | 1930 (77.0) |  |
| **At least one smoker family member** | **Present** | 492 (32.0) | 1538 (61.4) | <0.001 |
|  | **Absent** | 207 (21.4) | 967 (38.6) |  |
| **Smoker close friends** | **Some** | 472 (22.8) | 2074 (82.8) | <0.001 |
|  | **All** | 227 (32.5) | 431 (17.2) |  |
| **Curricula** | **4 years faculties** | 283 (20.8) | 1363 (54.4) | <0.001 |
|  | **2 years faculties** | 416 (36.4) | 1142 (45.6) |  |
| **Healthcare related faculty student** | **Yes** | 142 (22.4) | 634 (25.3) | <0.001 |
|  | **No** | 557 (29.8) | 1871 (74.7) |  |
| **Class** | 1 | 324 (29.0) | 1119 (44.7) | 0.193 |
|  | 2 | 251 (28.3) | 888 (35.4) |  |
|  | 3 | 55 (22.2) | 248 (9.9) |  |
|  | 4 | 69 (27.6) | 250 (10.0) |  |
| **Family type** | **Intact** | 32 (47.8) | 67 (2.7) | <0.001 |
|  | **Seperated** | 667 (27.4) | 2438 (97.3) |  |
| **Mother's education level** | **≤5 years** | 480 (26.3) | 1824 (72.8) | 0.004 |
|  | **>5 years** | 219 (32.2) | 681 (27.2) |  |
| **Father's education** | **≤5 years** | 297 (26.5) | 1120 (44.7) | 0.164 |
|  | **>5 years** | 402 (29.0) | 1385 (55.3) |  |
| **Smoking in the residency** | **Yes** | 315 (31.4) | 1004 (40.1) | 0.002 |
|  | **No** | 384 (25.6) | 1501 (59.9) |  |

**Table 2.** Sociodemographic characteristics of the students according to smoking status (Continued variables).

|  | Smoker (n:699)<br>mean±SD | Non smoker<br>(n:1806)<br>mean±SD | Total<br>(n:2505)<br>mean±SD |  |
| --- | --- | --- | --- | --- |
| **Age (min-max:17-46)** | 21.26±2.60 | 20.72±2.44 | 20.87±2.50 | <0.001 |
| **BMI (Kg/m<sup>2</sup>)** | 22.94±3.73 | 22.22±3.47 | 22.42±3.56 | <0.001 |
| **Number of family members** | 5.76±1.92 | 5.91±1.96 | 5.87±1.95 | 0.068 |
| **Number of siblings** | 3.56±1.74 | 3.80±1.85 | 3.73±1.83 | 0.003 |

Comparison of categorical demographic characteristics according to smoking status showed that the smoking rate was higher in the following populations: males compared to females, alcohol consumers compared to non-users, having all best friends who are smokers compared to some, having at least one smoker family member compared to none, studying in two-year faculties compared to four-year faculties, being in healthcare related faculties compared to other faculties, having destroyed/seperated families compared to intact families, having an at least secondary school graduated mother compared to lower levels, presence of smoking in the residency compared to absence (p<0.05) (Table 1).

Comparison of continuous variables showed thatcompared to non-smokers, smokers’ mean age was higher, mean BMI was higher, mean family member number was lower and accordingly mean number of siblings was lower (p<0.05) (Table 2).

Among smoker students, mean Fagerström test score was 4.74±1.16 for females and 5.09±1.31 for males (p<0.05); mean age of starting smoking was lower in males compared to females (Figure 1). Additionally males smoked 11 to 20 cigarettes per day on average, while females smoked less than 11 cigarettes daily (Figure 2). There was a negative correlation between nicotine dependence level and beginning age of smoking (r: -0.13, p<0.001) (Figure 3). Mean age for starting smoking was 16.34±2.72, for males 15.65±2.67, for females 17.03±2.60 (p<0.05) (Figure 2).

**Figure 1.**
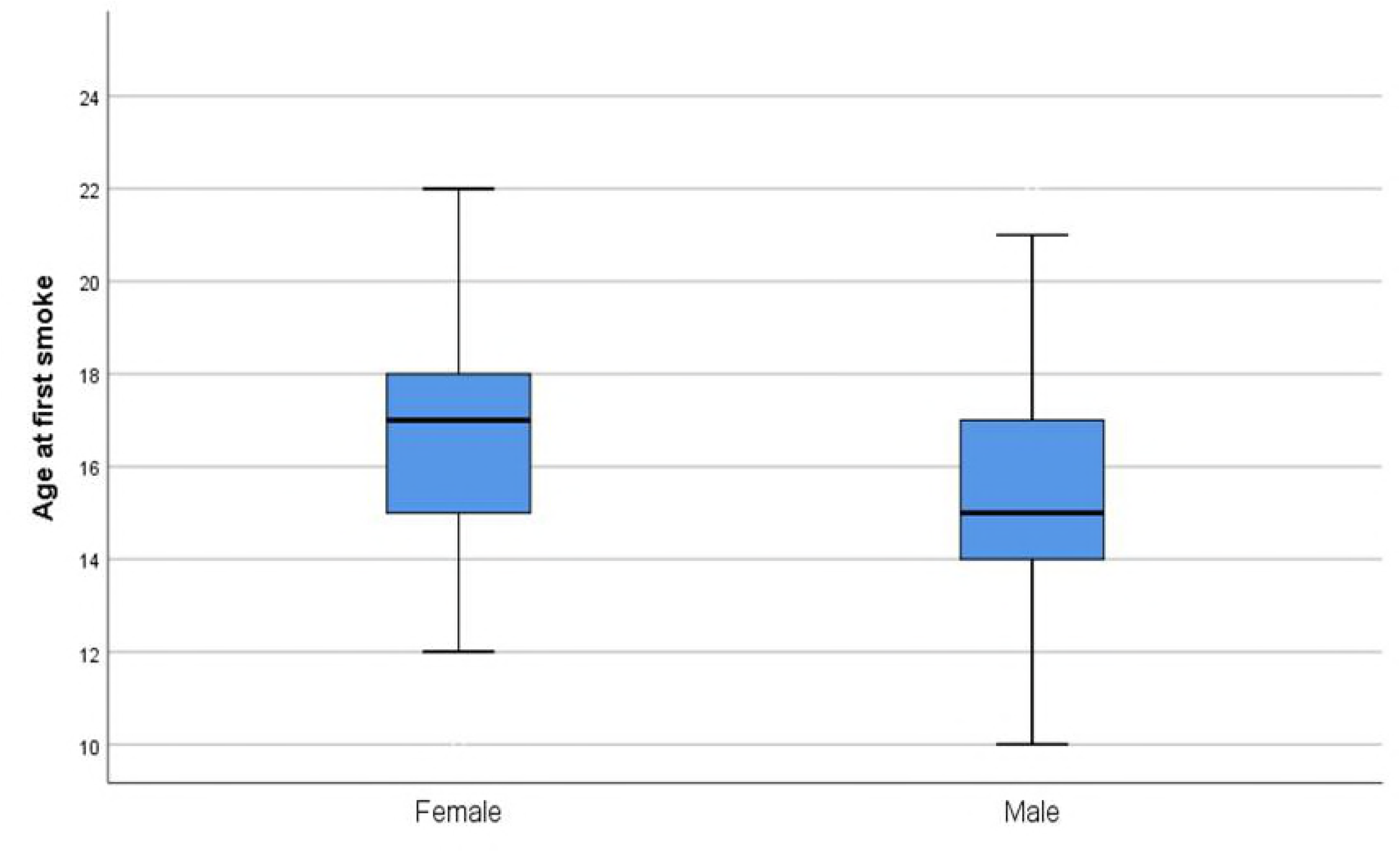
Age distribution at students’ first smoking experience according to gender.

**Figure 2.**
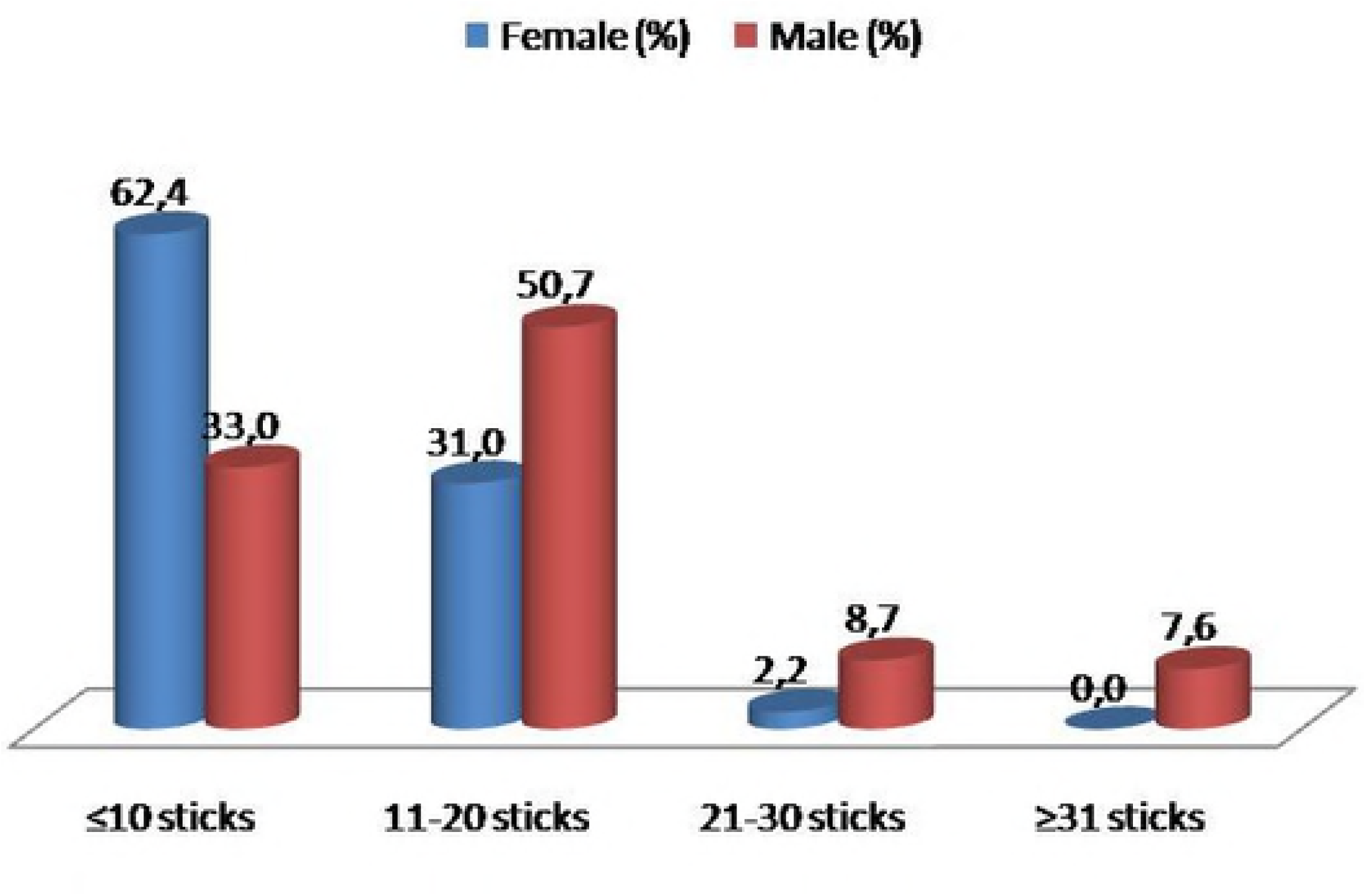
Daily smoked cigarette amount of students according to gender.

**Figure 3.**
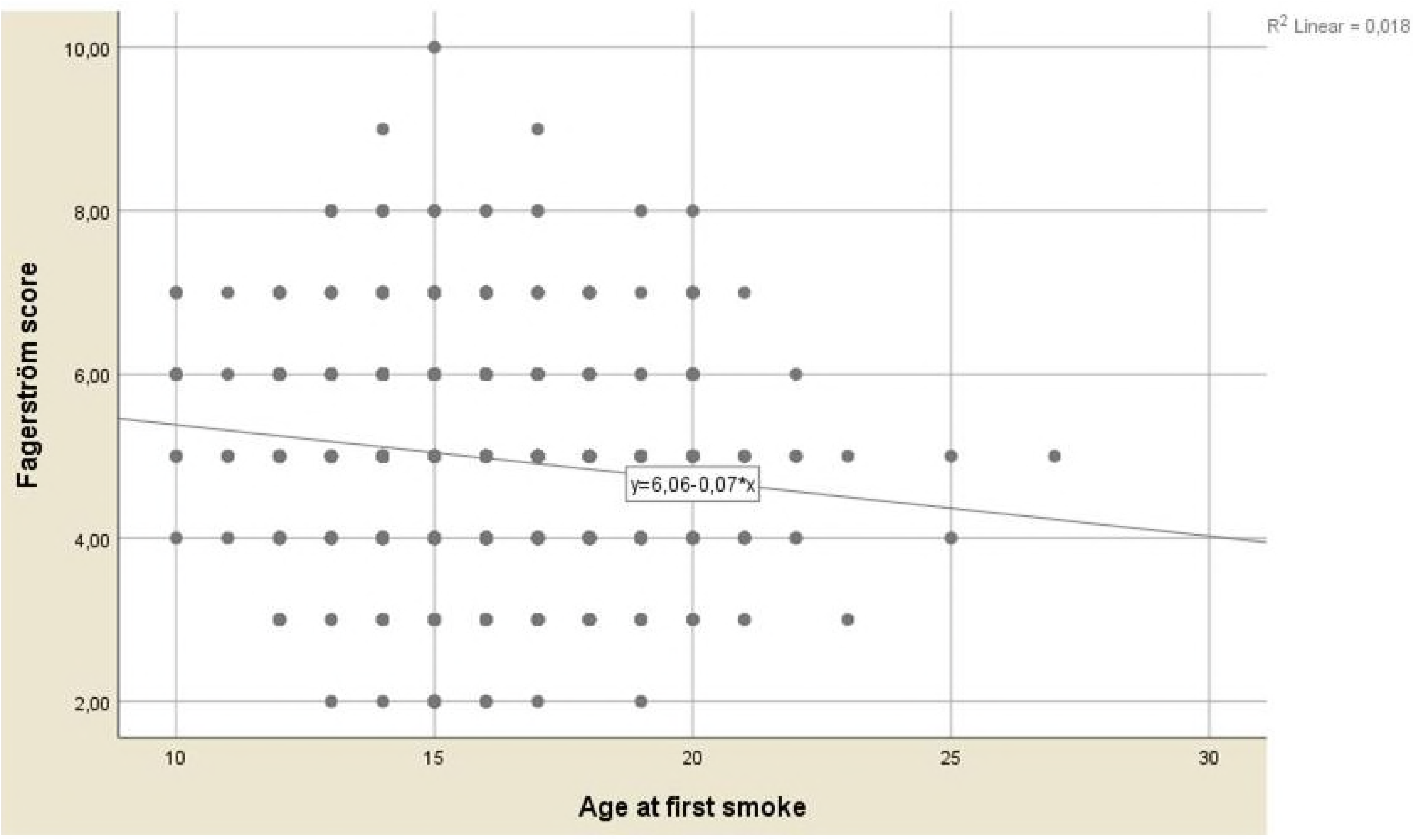
Correlation between the age of first smoke and the Fagerström score among current smokers

When the effect of these variables on the entire study population was evaluated through multivariate logistic regression analysis, it was seen that male gender (OR:3.43; 95%, CI:2.75-4.28), studying in two-year faculties (OR: 1.74; 95% CI: 1.39-2.18), having at least one close family member who is a smoker (OR:1.63; 95% CI:1.31-2.04), having all close friends who are smokers (OR:1.81; 95% CI: 1.40-2.33), and alcohol consumption (OR:4.39; 95% CI: 3.51-5.49) were positively associated with current smoking (p<0.05) (Table 3).

**Table 3.** Factors associated with smoking status in univariate and multivariate analysis.

|  | Univariate analysis |  |  | Multivariate analysis<br>(Backward: LR) |  |  |
| --- | --- | --- | --- | --- | --- | --- |
|  | OR | CI | p | OR | CI | p |
| Age (Per 1 age increment) | 1.083 | 1.047-1.120 | <0.001 | 1.037 | 0.995-1.082 | 0.088 |
| Gender (man v.s female) | 4.705 | 3.901-5.675 | <0.001 | 3.438 | 2.759-4.283 | <0.001 |
| Curriculae (2 years faculty vs. 4 years faculty) | 2.187 | 1.830-2.613 | <0.001 | 1.745 | 1.393-2.186 | <0.001 |
| Faculty (other than healthcare vs. healthcare) | 1.469 | 1.189-1.814 | <0.001 | 1.150 | 0.901-1.467 | 0.262 |
| BMI (Per 1 level increment) | 1.056 | 1.031- 1.082 | <0.001 | 0.988 | 0.959- 1.018 | 0.415 |
| Academic level (Per 1 level increment) | 0.942 | 0.859-1.033 | 0.203 | 0.989 | 0.871-1.122 | 0.863 |
| Family type (Destroyed vs. intact) | 2.428 | 1.491-3.953 | <0.001 | 1.543 | 0.843-2.827 | 0.160 |
| Number of family member (Per 1 level increment) | 0.958 | 0.915-1.003 | 0.068 | 1.008 | 0.954-1.064 | 0.782 |
| Mother's education level (>5 years vs. ≤5 years) | 1.117 | 1.047-1.192 | 0.001 | 1.050 | 0.817-1.351 | 0.701 |
| Father's education level (>5 years vs. ≤5 years) | 1.082 | 1.016-1.153 | 0.014 | 1.123 | 0.894-1.410 | 0.320 |
| Smoking in the residency (Present vs. absent) | 1.330 | 1.115-1.587 | 0.002 | 1.206 | 0.976-1.491 | 0.083 |
| At least one smoker family member (Presence vs. absence) | 1.727 | 1.432-2.083 | <0.001 | 1.639 | 1.314-2.044 | <0.001 |
| Smoker close friends (all vs. some) | 3.777 | 3.046-4.683 | <0.001 | 1.810 | 1.405-2.333 | <0.001 |
| Alcohol consumption (Presence vs. absence) | 6.537 | 5.339-8.004 | <0.001 | 4.391 | 3.511-5.491 | <0.001 |

Note: 2 Loglikelihood: 2344.930, R<sup>2</sup>: 0.220 (Cox&Snell), 0.316 (Nagelkerke), omnibustest of MCchi-square: 621.221, p=0.000,

## Discussion

This study looked into the prevalence of cigarette smoking and the associated factors that would affect smoking behavior among students at Çoruh University. The overall current smoking rate was 27.9%; 46% among males, 15.3% among females. This study showed that male gender, studying in two-year faculties, having exclusively best/close friends who are smokers, having at least one smoker family member, and consuming alcohol were positively associated with increased current smoking risks. Additionally, among daily smokers, male students had higher nicotine dependence levels and lower mean ages of smoking onset. A negative correlation was found between Fagerström test score and age at first cigarette.

According to data from Global Adult Tobacco Survey 2012, Turkey’s current smoking rate is 27.1%: 41.5 among males and 13.1 among females (20). According to the same survey, among the population ages 15-24 the current smoking rate is 33.0% in males and 7.4% in females. It is important to monitor the younger population’s smoking status periodically. This study was the first to evaluate the Çoruh University students’ smoking situation, therefore it is not possible to make a comparison against the smoking status of the university students in previous years. However it is possible to estimate general trends when comparing other universities’ results in recent years. According to current data, compared to the country’s overall cigarette smoking rate in that age group—and compared to university students’ smoking rates in studies conducted in last five years—we found our study population’s smoking levels to be higher for both genders (10, 20). Previous studies conducted among Turkish students revealed that living in urban areas was found to be a risk factor in their smoking habits (21). In our country, universities give young adults the opportunity to move away from their families, accordingly family pressure/control decreases and they start to spend most of their time with their peers. This may be one of the reasons for higher smoking rates among the students. Additionally, the location of our study, the city of Artvin, has a geographical border with Georgia and plays a significant role in the illicit tobacco transport from that country (16). That leads to the availability of cheaper cigarettes—almost half the price of those sold legally (7). It has been proven that increasing the cigarette tax discourages individuals from smoking (22, 23), therefore Turkey has followed MPOWER policies and increased the legal cigarette prices (3). However, in eastern regions of the country particularly illicit cigarettes and other types of tobacco products are prominent (7). The location of this study is nearby the illicit transport; this can be a factor in students’ ability to obtain cheap/illicit cigarettes with minimal effort. The high smoking rate in our study population can be further attributed to the ease of smoking and buying cigarettes on the university campus. In previous studies it was proven that smoking bans on university campuses decrease the smoking rate (12, 24); the university in the current study had no such ban. The lack of social and sport entertainment opportunities at the university may have also affected students’ smoking rates by contributing to student physical and mental inactivity. Therefore, to decrease smoking among this population and other university students, preventions such as smoking bans and offering student entertainment may be feasible and effective solutions.

Both univariate analysis and multivariate logistic regression analysis showed that males have higher smoking ratios (p<0.0001). This result is similar to other study findings both in Turkey (11) and globally (5, 18, 25). We concur with previous studies (5, 25) that this might have been because of the internalization of “gender roles” by the participants, and also because of the societal and cultural acceptance of the smoking habit among men rather than women. Also, female students’ daily cigarette use, and accordingly their nicotine dependence test results, were lower than males. One explanation for that distinction may be the higher mean age of women at the onset of smoking, because it has been previously reported that younger ages are associated with higher dependence and lower quit rates (26, 27). According to the Global Adult Tobacco Survey, in 2012 the mean age for starting smoking was 19;novel data shows that this age is decreasing. In our study it was found to be 16 on average (20). The overall smoking rate in this study was 27%, but the female students’ decreased rate had a significant effect on the average. Therefore it would be best in future studies to discuss the smoking rate according to gender instead of overall.

In our study and others, alcohol consumption increased the risk of smoking (14). Studies have shown that alcohol use is a risk factor for early initiation of cigarette smoking (28) and continuation of smoking (29). Additionally, we found that the students in two-year faculties had higher smoking rates. In Turkey the only difference between two– and four-year faculties is that acceptance into four-year faculties requires higher university exam marks. The effects of low academic success on smoking behavior have beenpreviously studied and low academic performance was found to be a predictor for smoking (30).

The strongest association in the current study was found between smoking and having both best friends and parents who smoke, which reflects results of previous studies (15, 18, 31). The influence of parental smoking seems less clear, but believing that family members smoke and having a positive attitude toward smoking were both factors that were predictive for smoking (32). In a recent study, living in intact families was shown to be a predictive factor for never smoking among students (33). In another study, living situation, mother’s educational level, economic status, and parents’ marital status were found to be the most influential predictive factors for substance abuse, including smoking (34). Studies have shown that overall smoking prevalence increases with age; similarly, the comparison of first and last year students’ smoking prevalence in Turkey shows that prevalence is higher among last year students compared to first (9, 35). In our results such factors had statistically significant effects on smoking status only in univariate analysis; at multivariate analysis these differences were not statistically significant.

### Strengths and limitations

This study was unique in several ways that made it valuable to the literature on Turkish smoking habits, particularly in the youth population. The high sample size of the study made it possible to generalize the results among university students in this region more easily. Similar, the researchers had the ability to evaluate not only four-year faculties but also two-year faculties—a gap in data that has been recorded as a need in previous studies (36). The location also allowed for data from a region that has a role in the illicit transport of tobacco products into the country. Furthermore, there have not been any studies on university students’ smoking behavior in the last 5 years, meaning that our study adds new informationto the data pool for this population. However, there are some limitations, including a lack of proof of illicit tobacco use and the study’s cross sectional design, which eliminates any temporal association. More studies from different regions of the country with both qualitative and quantitative measures are needed.

In conclusion, this study evaluated the smoking status and the factors affecting smoking habits of university students located in a small city inTurkey. Our results showed higher smoking rates among students compared to the country’s general population in that age range; the data was also higher compared to other Turkish university students according to recent data. That result may be related to not only the geographical location of the study but also the difference in academic success of the students. Even taking each of these factors into consideration, the smoking rate of our study population is alarming. It is important to review the government tobacco control policies in order to take new precautions for universities, including campus-wide smoking bans, increase in social/sport entertainment, and more educational activities about the harm of smoking.

## References

1. World Health Organization. WHO report on the global tobacco epidemic, 2017. Geneva, Switzerland: World Health Organization; 2017. Available from: http://www.who.int/tobacco/global_report/2017/en/

2. World Health Organization. The MPOWER Package. Geneva, Switzerland: World Health Organization; 2017. Available from: http://www.who.int/tobacco/mpower/en/

3. Elbek O, Kılınç O, Aytemur ZA, Akyildiz L, Küçük ÇU, Özge C, et al. Tobacco Control in Turkey. Turk Thorac J. 2015 Jul;16(3):141–150. doi:10.5152/ttd.2014.3898. Epub 2014 Jun 20.

4. Bilir N, Özcebe H. Tobacco control activities in Turkey. Turk J Public Health 2013; 11(2).

5. Wintemberg J, Yu M, Caman OK. Health Warnings, Smoking Rules, and Smoking Status: A Cross-National Comparison of Turkey and the United States. Subst Use Misuse. 2018 May 12;53(6):963–971. doi: 10.1080/10826084.2017.1387566.

6. Alvur TM, Cinar N, Oncel S, Akduran F, Dede C. Trends in smoking among university students between 2005-2012 in Sakarya, Turkey. Asian Pac J Cancer Prev. 2014;15(11):4575–81.

7. Kaplan B, Navas-Acien A, Cohen JE. The prevalence of illicit cigarette consumption and related factors in Turkey. Tob Control. 2017 Jun 30. pii: tobaccocontrol-2017-053669. doi: 10.1136/tobaccocontrol-2017-053669.

8. Centers for Disease Control and Prevention (CDC) Use of cigarettes and other tobacco products among students aged 13-15 years worldwide, 1999-2005. MMWR Morb Mortal Wkly Rep. 2006;55:553–6.

9. Özcebe H, Doğan BG, İnal E, Haznedaroğlu D, Bertan M. Smoking Habits and the Related Sociodemographic Characteristics in University Students. Turk Toraks Derg 2014; 15: 42–8.

10. Kilinç G, Bolgül BS, Aksoy G, Günay T. The Prevelance of Tobacco Use and the Factors Influencing in Students Studying at Two Dentistry Faculties in Turkey. Turk Thorac J. 2016 Apr;17(2):47–52. doi: 10.5578/ttj.17.2.010.

11. Şenol Y, Donmez L, Turkay M, Aktekin M. The incidence of smoking and risk factors for smoking initiation in medical faculty students: cohort study. BMC Public Health 2006, 6:128.

12. Gong M, Liang ZY, Zhang YY, Shadel WG, Zhou L, Xiao J. Implementation of the Tobacco-Free Campus Policy on College Campuses: Evidence From a Survey of College Students in Beijing. Nicotine Tob Res. 2016 Nov;18(11):2083–2091.

13. İçli F, Calişkan D, Gönüllü U, Sunguroğlu K, Akdur R, Akbulut H, et al. Fighting against cigarette smoking among medical students: a success story. J Cancer Educ. 2014 Sep;29(3):458–62. doi: 10.1007/s13187-013-0573-y.

14. Eticha T, Kidane F. The prevalence of and factors associated with current smoking among College of Health Sciences students, Mekelle University in northern Ethiopia. PLoS One. 2014 Oct 23;9(10):e111033. doi: 10.1371/journal.pone.0111033. eCollection 2014.

15. Metintaş S, Sariboyaci MA, Nuhoğlu S. Smoking patterns of university students in Eskişehir, Turkey. Public Health. 1998 Jul;112(4):261–4.

16. Illicit cigarette transport. Available from: http://www.artvm.pol.tr/Haberier/Sayfalar/ARTViN’DE-REKOR-KA#AK-SiGARA-YAKALAMASI.aspx

17. Çakmakçi Karadoğan D, Önal Ö, Say Şahin D, Yazıcı S, Kanbay Y. Evaluation of school teachers’ sociodemographic characteristics and quality of life according to their cigarette smoking status: a cross-sectional study from the eastern Black Sea region of Turkey. Tuberk Toraks. 2017 Mar;65(1):18–24.

18. Yurt Öncel S, Gebizlioğlu OL, Aliev Alioğlu F. Risk factors for smoking behavior among university students. Turk J Med Sci 2011; 41 (6): 1071–1080. doi:10.3906/sag-1009-1122.

19. Heatherton TF, Kozlowski LT, Frecker RC, Fagerström KO. The Fagerström Test for Nicotine Dependence: a revision of the Fagerström Tolerance Questionnaire. Br J Addict. 1991 Sep;86(9):1119–27.

20. Public Health Institution of Turkey. Global Adult Tobacco Survey Turkey 2012. Available from: http://www.who.int/tobacco/surveillance/survey/gats/report_tur_2012.pdf

21. Akca G, Guner SN, Akca U, Kilic M, Sancak R, Ozturk F. Students’ unchanging smoking habits in urban and rural areas in the last 15 years. Pediatr Int. 2016 Apr;58(4):279–83. doi: 10.1111/ped.12814.

22. Burki TK. WHO tobacco report focuses on increased taxation. Lancet Respir Med. 2015 Aug;3(8):604. doi: 10.1016/S2213-2600(15)00292-1. Epub 2015 Jul 22. PMID: 26210312 DOI: 10.1016/S2213-2600(15)00292-1.

23. Kostova D, Andes L, Erguder T, Yurekli A, Keskinkiliç B, Polat S, et al. Cigarette prices and smoking prevalence after a tobacco tax increase‐‐Turkey, 2008 and 2012. MMWR Morb Mortal Wkly Rep. 2014 May 30;63(21):457–61.

24. Erdal G, Erdal H, Esengun K, Karakas G. Cigarette consumption habits and related factors among college students in Turkey: a logit model analysis. J Pak Med Assoc. 2015 Feb;65(2):136–41.

25. Mbatchou Ngahane BH, Luma H, Mapoure YN, Fotso ZM, Afane Ze E. Correlates of cigarette smoking among university students in Cameroon. Int J Tuberc Lung Dis. 2013 Feb;17(2):270–4. doi: 10.5588/ijtld.12.0377.

26. Khuder SA, Dayal HH, Mutgi AB. Age at smoking onset and its effect on smoking cessation. Addict Behav. 1999 Sep-Oct;24(5):673–7.

27. Darville A, Hahn EJ. Hardcore smokers: what do we know? Addict Behav. 2014 Dec;39(12):1706–12. doi: 10.1016/j.addbeh.2014.07.020.

28. So ES, Yeo JY. Factors Associated with Early Smoking Initiation among Korean Adolescents. Asian Nurs Res (Korean Soc Nurs Sci). 2015 Jun;9(2):115–9. doi: 10.1016/j.anr.2015.05.002.

29. Kim H, Kim MH, Park YS, Shin JY, Song YM. Factors That Predict Persistent Smoking of Cancer Survivors. J Korean Med Sci. 2015 Jul;30(7):853–9. doi: 10.3346/jkms.2015.30.7.853. Epub 2015 Jun 10.

30. Kinnunen JM, Lindfors P, Rimpelä A, Salmela-Aro K, Rathmann K, Perelman J, et al. Academic well-being and smoking among 14- to 17-year-old schoolchildren in six European cities. J Adolesc. 2016 Jul;50:56–64. doi: 10.1016/j.adolescence.2016.04.007.

31. Mak KK, Ho SY, Day JR. Smoking of parents and best friend‐‐independent and combined effects on adolescent smoking and intention to initiate and quit smoking. Nicotine Tob Res. 2012 Sep;14(9):1057–64. doi: 10.1093/ntr/nts008.

32. Escario JJ, Wilkinson AV. The Intergenerational Transmission of Smoking Across Three Cohabitant Generations: A Count Data Approach. J Community Health. 2015 Mar 22.

33. Du Y, Palmer PH, Sakuma KL, Blake J, Johnson CA. The Association between Family Structure and Adolescent Smoking among Multicultural Students in Hawaii. Prev Med Rep. 2015;2:206–212.

34. Jalilian F, Karami Matin B, Ahmadpanah M, Ataee M, Ahmadi Jouybari T, Eslami AA, et al. Socio-demographic characteristics associated with cigarettes smoking, drug abuse and alcohol drinking among male medical university students in Iran. J Res Health Sci. 2015 Winter;15(1):42–6.

35. Kocabas A, Burgut R, Bozdemir N, Akkoçlu A, Çildağ O, Dağlı E, et al. Smoking patterns at different medical schools in Turkey. Tob. Control 1994;3(3):228–235.

36. Bennett BL, Deiner M, Pokhrel P. College anti-smoking policies and student smoking behavior: a review of the literature. Tob Induc Dis. 2017 Feb 1;15:11. doi: 10.1186/s12971-017-0117-z.

